# Human IgG neutralizing monoclonal antibodies block SARS-CoV-2 infection

**DOI:** 10.1101/2020.05.19.104117

**Authors:** Jinkai Wan, Shenghui Xing, Longfei Ding, Yongheng Wang, Dandan Zhu, Bowen Rong, Siqing Wang, Kun Chen, Chenxi He, Songhua Yuan, Chengli Qiu, Chen Zhao, Xiaoyan Zhang, Xiangxi Wang, Yanan Lu, Jianqing Xu, Fei Lan

## Abstract

The coronavirus induced disease 19 (COVID-19) caused by the severe acute respiratory syndrome coronavirus 2 (SARS-CoV-2) has become a worldwide threat to human lives, and neutralizing antibodies present a great therapeutic potential in curing affected patients. We purified more than one thousand memory B cells specific to SARS-CoV-2 S1 or RBD (receptor binding domain) antigens from 11 convalescent COVID-19 patients, and a total of 729 naturally paired heavy and light chain fragments were obtained by single B cell cloning technology. Among these, 178 recombinant monoclonal antibodies were tested positive for antigen binding, and the top 13 binders with K_d_ below 0.5 nM are all RBD binders. Importantly, all these 13 antibodies could block pseudoviral entry into HEK293T cells overexpressing ACE2, with the best ones showing IC50s around 2-3 nM. We further identified 8 neutralizing antibodies against authentic virus with IC50s within 10 nM. Among these, 414-1 blocked authentic viral entry at IC50 of 1.75 nM and in combination with 105-38 could achieve IC50 as low as 0.45 nM. Meanwhile, we also found that 3 antibodies could cross-react with the SARS-CoV spike protein. Altogether, our study provided a panel of potent human neutralizing antibodies for COVID19 as therapeutics candidates for further development.

## Introduction

Over the last two decades in 21th century, the outbreaks of several viral infectious diseases affected millions of people^1–7^. Among these, 3 coronaviruses, SARS-CoV, MERS and SARS-CoV-2^8^, have received significant attention especially due to current outbreak of COVID-19 caused by SARS-CoV-2, and high mortality rates of the infected individuals. Most patients died due to severe pneumonia and multi-organ failure^4,9^. Despite of rare exceptions, such as asymptomatic carriers, exist, it is generally believed if the infected individuals could not develop effective adaptive immune responses for viral clearance to prevent sustained infection, there are high chances for transformation into severe acute respiratory infection. Supporting this idea, treatment with convalescent plasma to COVID-19 patients showed significant clinical improvement and decreased viral load within days^10^. However, the sources of convalescent plasma are limited and could not be amplified, therefore, effective and scalable treatments are still urgently needed^11^.

Owing to recent rapid development of single cell cloning technology, the process of antibody identification has been much shortened, from years to even less than 1 month. Therefore, full human and humanized neutralizing antibodies represent as great hopes for a prompt development of therapeutics in treating infectious diseases. In support of this, cocktail treatment of 3 mixed antibodies recognizing different epitopes, with one of them able to robustly neutralize, was successfully used in the curation of a British Ebola patient^12,13^. Regarding coronaviruses, neutralizing antibodies against MERS were tested effective in animals^14^. While SARS-CoV neutralizing antibodies did not meet human due to lack of patients after the development. Interestingly one of them, S309, was shown to be able to cross-react with and neutralize SARS-CoV-2^15^. However, the RBD regions (key targets for viral neutralization, also see below) only share 74% sequence identity between the two SARS viruses^16^, raising concerns about the effectiveness of SARS-CoV neutralizing antibodies against SARS-CoV-2.

The spike proteins of coronaviruses play an essential role in viral entry into human target cells. The S1 region, especially the RBD domain, primes the viral particle to human cell surface through the interaction with the receptor protein Angiotensin I Converting Enzyme 2 (ACE2)^17^, which then triggers infusion process primarily mediated by S2 region^18^. The primary amino acid sequences of the spike proteins of SARS-CoV and SARS-CoV-2 share 76 % identity throughout the full coding regions, with 79.59 % similarity and 74% identity in RBD domains^16,19,20^. Structure analyses revealed high 3D similarity between the spike proteins of the two viruses, and both trimerize and interact with ACE2 through the RBD domains^21^. Importantly, the interaction between the SARS-CoV-2 RBD and ACE2 was assessed at around 1.2 nM^18,22^, 4 folds stronger than the SARS-CoV RBD. While this enhanced affinity may explain a much stronger spreading ability of SARS-CoV-2, it also suggests that finding potent neutralizing antibodies targeting SARS-CoV-2 RBD could also be more challenging.

While this manuscript was under preparation, identification of multiple human neutralizing monoclonal antibodies had been reported by a few studies^23–27^. While these groups and ours all employed similar approaches and obtained authentic viral neutralizing antibodies, the performances of these antibodies varied largely in different assays, e.g. the correlation between binding affinities, pseudoviral and authentic viral neutralizing abilities. Nevertheless, more human antibodies either directly neutralizing SARS-CoV-2 or opsonizing free viral particles for rapid immune clearances are still needed.

Here, we report the identification of 178 S1 and RBD binding full human monoclonal antibodies from the memory B cells of 11 recently recovered patients. The CDR3 sequences of the vast majority of these antibodies are different, indicating they are developed from different B cell clones. A total of 13 antibodies showed binding affinities lower than 0.5 nM and pseudoviral neutralizing abilities. We further identified 8 antibodies showing robust authentic viral neutralizing capabilities. Among these, the best one, 414-1, showed authentic viral neutralization IC50 at 1.75 nM. Moreover, we also identified 3 antibodies cross-reacting with SARS-CoV spike protein.

## Results

### Serological responses and single B cell isolation

We screened 11 patients recently recovered from COVID-19, and identified 9 out of 11 individuals with strong serological responses to SARS-CoV-2 Spike RBD and S1 protein (Figure 1 A and 1B), and 7 sera showed neutralization abilities for SARS-CoV-2 pseudoviral infection of HEK293T cells stably expressing human ACE2 (Figure 1B). Therefore, sera from different individuals displayed a wide range of antibody responses to SARS-CoV-2 infection in our assays, consistent with a recent report^28^.

**Fig.1.**
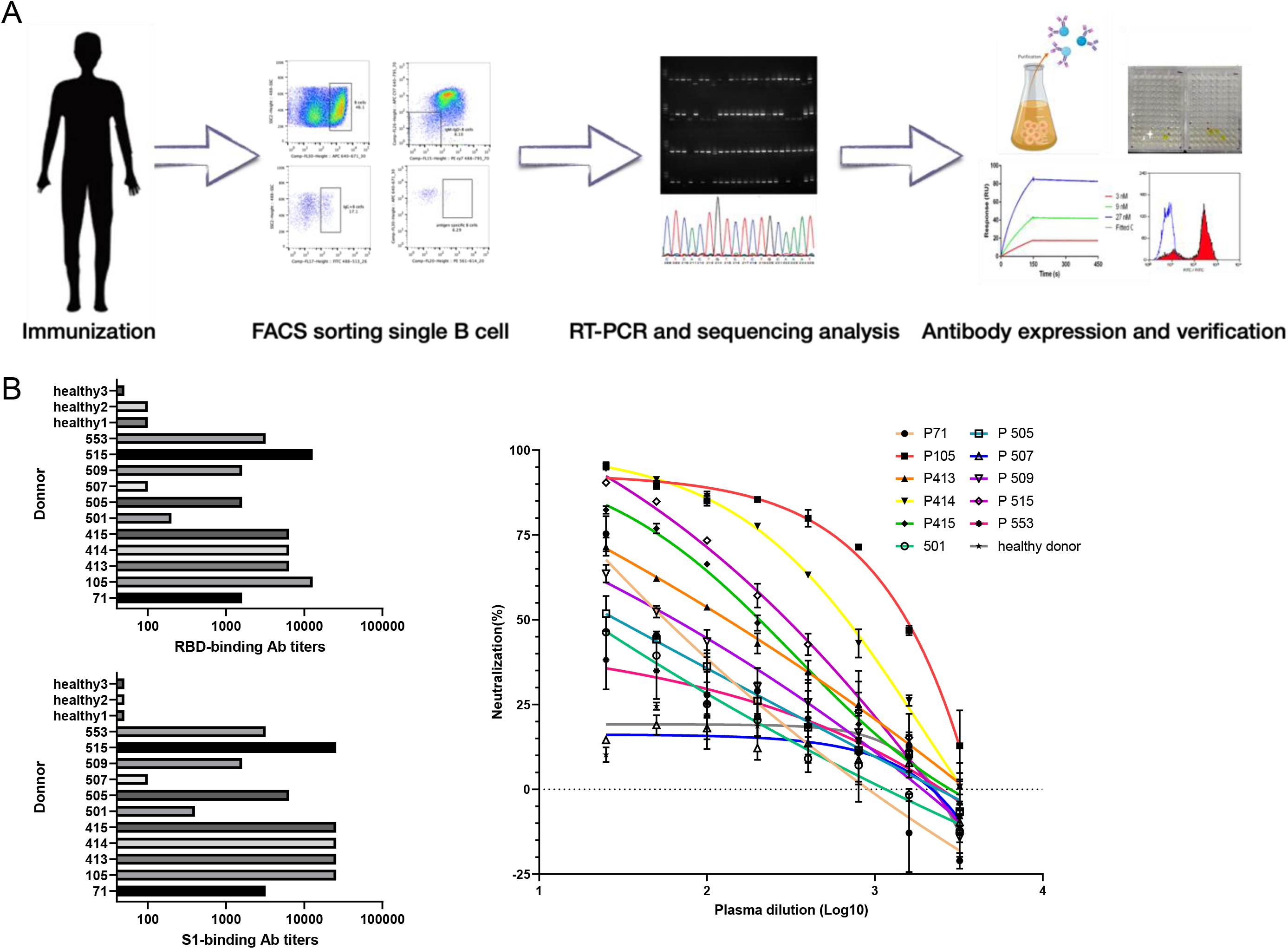
Single B cell isolation strategy and serological responses of 11 convalescent COVID19 patients. (A) Schematic depicting the screening strategy used to sort B cells from SARS-CoV-2 convalescent patients and antibody identification. (B) Spike protein binding and pseudoviral neutralizing tests of donor plasma. RBD and S1 were used. Plasma samples of heathy donors were used as controls. The mean values and standard deviations of two technical replicates were shown.

The RBD domain in the S1 region of SARS-CoV-2 spike protein is the critical region mediating viral entry through host receptor ACE2. Using recombinant viral antigens, we then isolated RBD and S1 binding memory B cells for antibody identification using the PBMCs (peripheral blood mononuclear cells) from 11 individuals by flow cytometry-based sorting technology (Figure 1A). Each individual exhibited different frequencies of viral antigen specific memory B cells (Figure S1 and Table S1).

Sequences encoding antibody heavy (IGH) and light (IGL) chains were amplified from single B cell complementary DNA samples after reverse transcription and then cloned through homologous recombination into mammalian expressing vectors^29^. Overall, 729 naturally paired antibody genes were obtained from the 11 individuals, of which the No.71 individual failed to give any positive antibody. However, no strong correlation was found between serological responses and the number of acquired SARS-CoV-2 S-specific antibodies (Figure 1B and Table S1), while we could not rule out possibilities of certain technical variations being involved. Of note, No.509 blood sample was obtained at the second day after hospitalization (Table S2), the sera already showed weak S-specific affinity and pseudoviral neutralizing capacity, despite that no strong antibodies were obtained from No.509 sample.

### Identification and affinity characterization of human monoclonal antibodies to SARS-CoV-2

All the 729 antibodies were further expressed in HEK293E cells and the supernatants were tested in ELISA for S1 or RBD binding (Figure 2A and Figure S2). Among these, 178 antibody supernatants were positive for RBD or S1 binding. We then purified all these antibodies in larger quantities to measure the precise values of K_d_ (EC50), and found the values varied broadly, with the most potent one at 57 pM (8.55 pg/μl), and 13 strongest ones having K_d_ below 0.5 nM (Figure 2B). All the positive clones were then sequenced. Notably, almost all (98.6%) of the sequences obtained were unique ones (Figure 2C, left), unlike what were previously reported for HIV-1, influenza and ZIKV^29^. Subsequently, we aligned the CDR3 of heavy chain (CDR3H) sequences of all these 13 antibodies and found 11 sequences are different, with the exceptions of 515-1 and 505-5 (Figure 2C, right). Interestingly, 515-1 and 505-5 also shared the same CDR3_L_ sequences, and they indeed behaved very similarly in the subsequent tests (see below).

**Fig.2.**
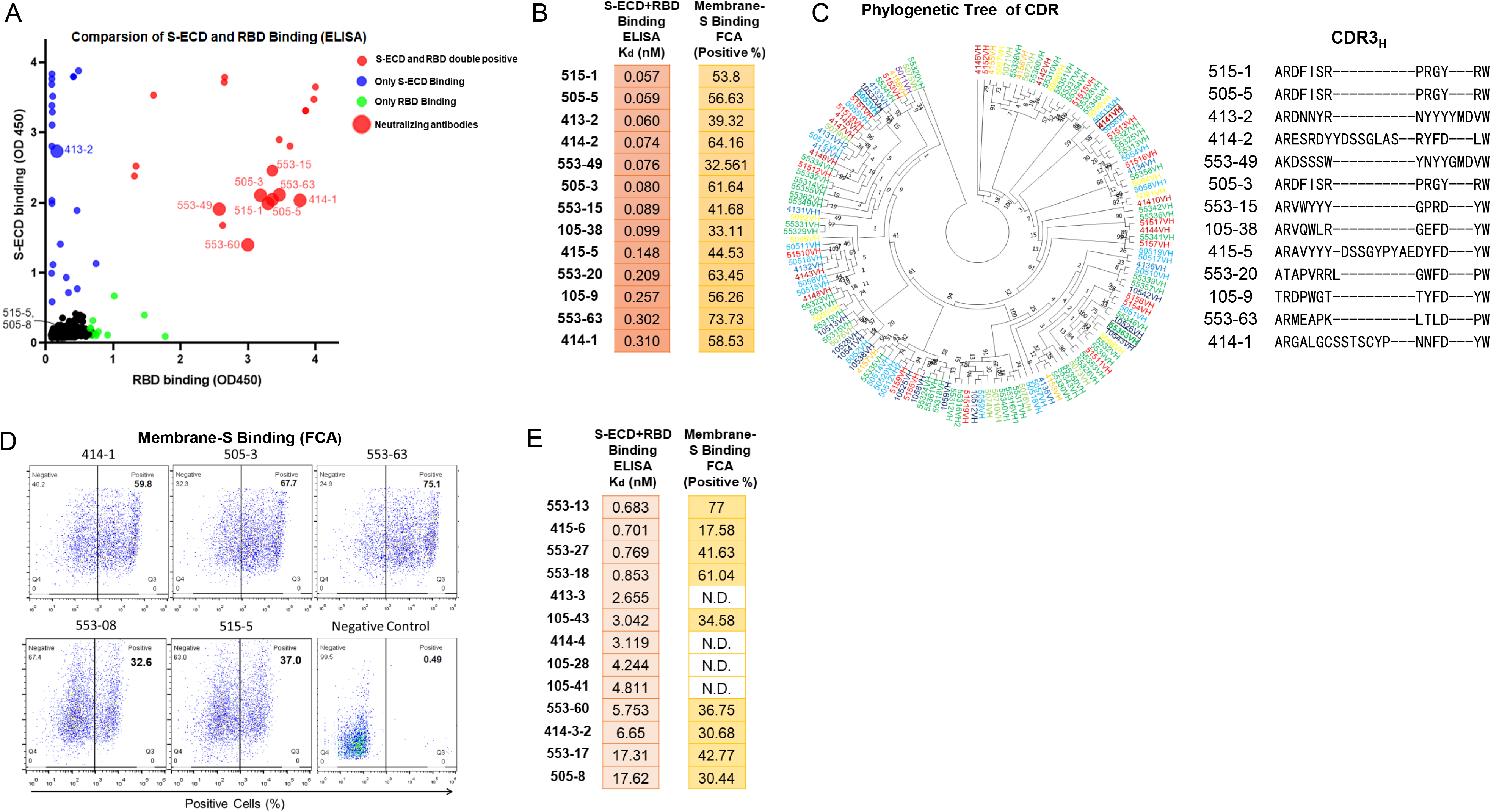
The identification and characterization of spike protein-specific monoclonal antibodies. (A) Characteristics of antibodies binding with RBD and S-ECD. RBD and S-ECD dual binders are presented in red dots, binders for S-ECD only are in purple, and binders for RBD only are in green. Authentic neutralizing antibodies are represented by bigger dots. (B) The flow cytometry binding results for the 13 strongest antibodies (K_d_ below 0.5 nM) were shown. (C) Left, Maximum-likelihood phylogenetic tree analysis of the heavy chains of all sequenced monoclonal antibodies. Different colors indicate antibodies identified from individual patients. Right, the alignment of the 13 CDR3_H_ sequences. (D) Flow cytometry analysis of representative antibodies binding to SARS-CoV-2 S protein expressing on A549 cell membrane. IgG-Fc-PE antibody was used as control. (E) Flow cytometry binding results of 13 less strong antibodies (K_d_ between 0.5 to 20 nM) were shown. N.D., non-detectable.

As the spike S1 protein of SARS-CoV-2 tends to undergo conformation changes during storage, we also performed flow cytometry analyses for all 729 purified antibodies or supernatants for the binding ability of the freshly expressed spike protein in the membrane bound form using HEK293T cells (Figure 2D). Among these 729 antibodies, 58 were obtained from B cells purified by recombinant RBD domain, and 671 were from B cells purified by recombinant S1 protein. From the latter, 135 binders were identified, and 21 of them were able to bind S-ECD (S protein extracellular domain) while showing low or no RBD affinity, tested by ELISA. The result indicated that RBD regions are the primary antigen inducing antibody generation and recognition (Figure 2A).

We also found that the results of the flow cytometry and ELISA assays were largely consist among the top 13 antibodies (K_d_ < 0.5 nM) detected by ELISA, all 13 were RBD binders and could recognize SARS-CoV-2 Spike protein on cell membrane by flow cytometry (Figure 2B). However, 4 (namely 413-3, 414-4, 105-28 and 105-41) of 13 less strong antibodies examined by ELISA (K_d_ between 0.5-20 nM) could not bind cell membrane S protein (Figure 2E). On the other hand, 2 antibodies 505-17 and 515-15, barely showing ELISA signals displayed strong affinities of S protein expressing on A549 cell membrane detected by flow cytometry (Figure S3). These observations indicated that the membrane-bound and soluble recombinant S proteins might have certain conformation alterations.

### Identification of potent neutralizing antibodies by pseudoviral and live viral infection assays

To identify neutralizing antibodies, we first employed pseudoviral infection assays using HEK293T-ACE2 cells. From all the antibodies tested, we found a total of 15 pseudoviral neutralizing antibodies. The best 3 antibodies in neutralizing pseudoviruses are 414-1, 505-3 and 553-63 (Figure 3 A and 3B, IC50 from 2.3 to 3.6 nM), all of which showed strong affinities towards RBD domain (Figure 3A and 3B, K_d_ from 0.079 nM to 0.31 nM) and could also robustly block RBD and ACE2 binding using ELISA (Figure 3C), indicating certain level of correlation between these abilities. To our surprise, two of the 15 pseudoviral neutralizing antibodies, 413-2 and 505-8, only bound RBD very weakly in ELISA (Table S3), but were able to strongly recognize membrane-S protein overexpressing in HEK293T cells (Table S3), indicating there is an alternative neutralization mechanism of non-RBD binders.

**Fig.3.**
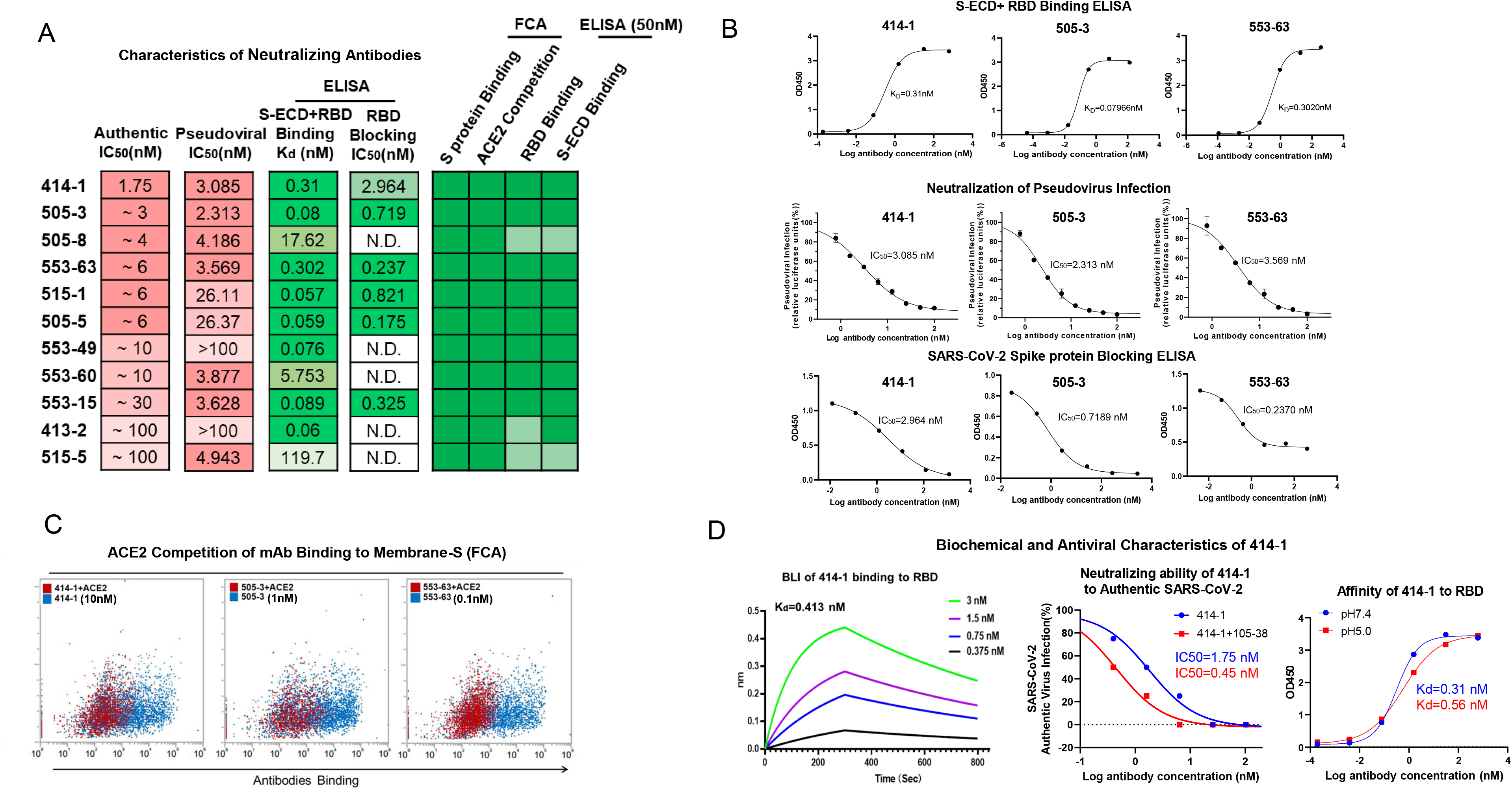
Neutralizing capacities of the leading monoclonal antibodies against pseudotyped and authentic viruses of SARS-CoV-2. (A) Summary of the indicated characteristics of all neutralizing antibodies. Red highlights antibodies with good characteristics, and green represents strong binding abilities (by flow cytometry) towards Spike protein expressed on A549 cell membrane. (B) Pseudoviral neutralizing and ACE2 blocking ELISA analyses of 414-1, 505-3 and 553-63. (C) ACE2 competition results of 414-1, 505-3 and 553-63 using flow cytometry. The blue dots represent A549-Spike treated with antibodies only, and the red dots represent A549-Spike incubated with 50nM human ACE2 together with 0.1, 1 and 10 nM antibodies, respectively. (D) Left, BLI examination of 414-1 and RBD binding affinity; Middle, neutralization results of 414-1 and 414-1 in combination with 105-38 against authentic virus (SARS-CoV-2-SH01) using Vero-E6; Right, 414-1 affinity examination by ELISA at pH 5 and 7.4.

Among the 15 pseudoviral neutralization antibodies, all could bind S protein expressed in cell membrane detected by flow cytometry analyses. To further validate these antibodies, we employed two ACE2 competition assays. One assay was to test the ability in blocking ACE2 and RBD binding using ELISA, and the second assay was to compete free ACE2 binding to S protein expressing in A549 membrane using flow cytometry (Figure 3C and Table S3). In order to perform the latter assay, we first tested various concentrations of soluble PE-labeled ACE2 in labeling 10 thousands of A549 overexpressing S protein, and found 50 nM ACE2 could label the vast majority of the cells (Figure S3). Then, we used 0.1, 1 and 10 nM antibodies to compete 50 nM ACE2 for the binding of S on A549 membrane. All 15 pseudoviral neutralizing antibodies, except for 553-20, could compete ACE2 (Figure S4A). However, the competition abilities did not show significant correlation with neutralizing abilities. For instances, 414-1 and 553-63 are the strongest neutralizing antibodies, however, 0.1 nM 553-63 could reach 50% competition against 50 nM ACE2 while 414-1 needed 10 nM (Figure 3C and Figure S4B).

We then performed authentic viral neutralization assay for the best 11 antibodies using Vero-E6 (Figure 3A), and found a total of 8 antibodies showing IC50 at for below 10 nM. The best one, 414-1, was able to effectively block authentic viral entry at IC50 of 1.75 nM, and when combined with 105-38 the IC50 could further reduced to 0.45 nM (Figure 3D). To note, although 105-38 showed a much weaker neutralizing ability by itself in pseudoviral assay, but it recognized different epitope as 414-1 (data not shown), explaining the combinatorial enhancement. We also tested 414-1 expressing in CHO cells, and found it could achieve ~ 300 mg/L without any optimization suggesting for great potential in therapeutic development. Subsequently, 414-1 was also tested in pH 5.0 for affinity maintenance by ELISA (Figure 3D), the results (K_d_= 0.31 nM in pH7.4 and K_d_= 0.56 nM in pH5.0) showed a good affinity performance in low pH, indicating less probability in causing ADE. Furthermore, 414-1 was also tested by BLI, and showed comparable K_d_ (Figure 3D).

### Cross-reactivity with SARS-CoV Spike protein

The spike proteins of SARS-CoV-2 share 76% and 35% of amino acid identities with SARS-CoV and MERS-CoV, respectively. Therefore, we next tested whether our antibodies could cross-react with the S proteins of these two other coronaviruses. In order to do so, we overexpressed the S proteins of SARS-CoV-2, SARS-CoV and MERS-CoV in HEK293T, and tested the cross-reactivities by flow cytometry analyses. After removal of the antibodies showing non-specific binding to HEK293T cells, we focused on 30 antibodies with robust membrane SARS-CoV-2 S protein binding abilities (Figure 4A), among them, we found 3 antibodies, 415-5, 415-6 and 515-5, recognizing SARS-CoV S, but none could recognize MERS-CoV S (Figure 4B). 415-5 and 515-5 shared similar S protein affinities between SARS-CoV-2 and SARS-CoV, but 415-6 had much lower affinity towards SARS-CoV S (Figure 4C).

**Fig.4.**
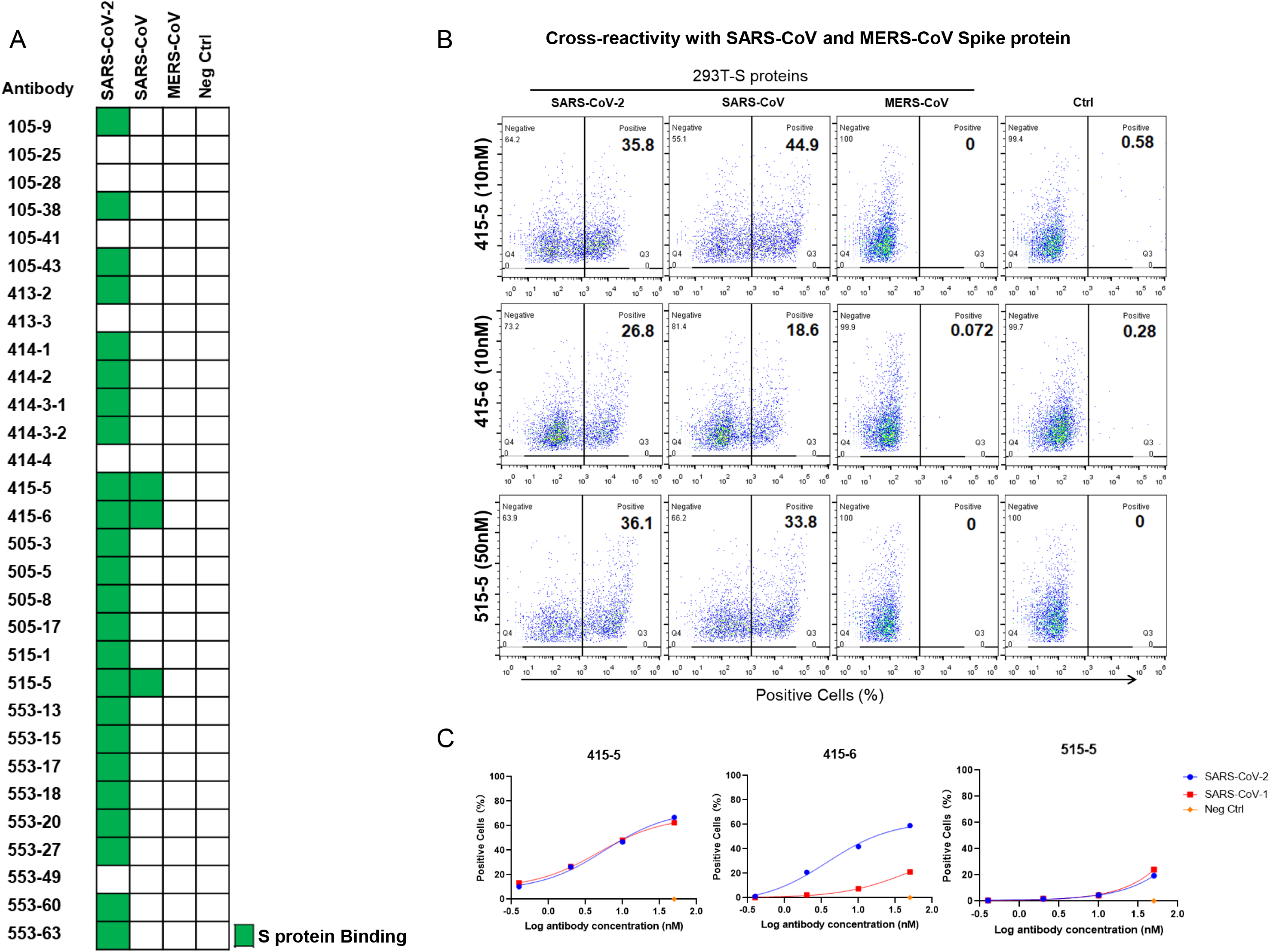
Cross-reactivity with SARS-CoV. Cross-reactivity analysis of the representative antibodies against S proteins of SARS-CoV and MERS-CoV. Flow cytometry analyse were performed using HEK293T cells expressing the S proteins of SARS-CoV-2, SARS-CoV and MERS-CoV, and 50 nM of the indicated antibodies were used. Non-transfected HEK293T were used as controls. (A) Heatmap summary of the results. (B-C) Flow cytometry data of the 3 cross-reactive antibodies.

## Discussion

Within last few weeks, several groups had published a panel of full human neutralizing antibodies against COVID19 ^23–27,30^. To date, a total of 28 human antibodies had been reported to show neutralizing abilities against authentic SARS-CoV-2, and here we added 11 more to the reservoir. Among these, 20 reported ones showed neutralizing IC50 within 10 nM, a general potency guideline for potential clinical use, and here we reported 8 new ones.

In our study, in order to provide good therapeutic candidates of monoclonal antibodies, we have implemented multiple approaches during the triage processes before authentic viral testing. Our triage approaches included, 1) RBD and S binding abilities by ELISA; 2) membrane S binding abilities by flow cytometry; 3) ACE2 competition abilities using both ELISA and flow cytometry; 4) pseudoviral neutralization assay. The best 3 antibodies, 414-1, 505-3 and 553-63, showed consistent performances in all these assays (Table S3). The other 5 antibodies with IC50 within 10nM for authentic viral neutralization also showed relatively consistent performances in most of the assays (Table S3). In general, the 7 of the top 8 neutralizing antibodies showed robust binding abilities in the ELISA assays. The only one exception is 505-8, however, it functioned well in flow cytometry-based binding and competition assays (Table S3, column 6-7, also see below), indicating that binding affinity is a fundamental quality for neutralizing capacity in our study. And we observed a general increase of neutralizing IC50 values compared to binding affinity EC50 values. However, compared with other studies mentioned above, we found that the leading antibodies reported in the studies by Cao et al^23^ and Yan et al^27^, showed great neutralization abilities, IC50s ranging from 0.1 - 1 nM, showed EC50s of 0.8 - 70 nM. Similarly, a 100-fold enhancement of neutralizing activity compared to the affinity was also reported by Chi et al for their leading antibodies^25^. The reason causing such discrepancy at this point is yet unknown, however, conformation alterations between recombinant S and RBD protein and their natural forms may be one possibility, and assay conditions could also vary among different groups.

Among the top 30 antibodies, 24 were obtained from B cells isolated using S1 protein. We found the majority, 18 of 24, are RBD binders, and two of them, 413-2 and 553-13, are strong S binders but not RBD binders tested by ELISA (Table S3, last two columns). These findings indicated that the RBD region is the primary epitope for immune responses. Similar observations were also reported elsewhere for SARS-CoV-2^23,27^ as well as SARS-CoV^31,32^. We also found another 4 antibodies, 505-8, 505-17, 515-5 and 553-18, showed barely any ELISA signals towards both RBD and S, but could robustly bind freshly expressed S protein in A549 membrane (Table S3). These included one neutralizing antibody 505-8 mentioned above, indicating that the recombinant RBD or S protein may differ from the membrane bound S in terms of 3D conformation. We would have missed these antibodies if we only had used ELISA for antibody triage. Therefore, future antibody study should consider multiple approaches for the initial identification and quality control.

It’s encouraging to see the increasing numbers of human neutralizing antibodies against COVID-19. The combination of different clones of potent neutralizing human monoclonal antibodies recognizing different vulnerable sites of SARS-CoV2 can prevent the occurrence of mutant escapes when administered clinically. Therefore, more candidate antibody sequences could greatly facilitate the therapeutic development in curing COVID-19 patients.

## Acknowledgments

We appreciate the Novoprotein Scientific Inc. for gifting SARS-CoV-2 Spike, ACE2 and related recombinant proteins. Prof. Lu Lu and Prof. Zhigang Lu from Fudan University provided 293T-ACE2 cell line and helpful suggestions. We thank Prof. Lilin Ye from Third Military Medical University for helpful suggestions and manuscript writing. We thank Yiwei Tang from Fudan University for helps in drawing the maximum-likelihood phylogenetic tree. HEK293E cell line was a gift from Prof. Yanhui Xu, Fudan University.

## Author contributions

F.L., J.X., Y.L. and X.W. conceived the project. J.W., S.X., L.D., Y.W. S.Y., D.Z., B.R., S.W., K.C., C.H., C.Z. and C.Q. did the experiments. All authors contributed to data analyses. F.L., J.W., S.X., L.D. and Y.L. wrote the manuscript.

## Materials and methods

### Ethics statement

The experiments involving authentic COVID-19 virus were performed in Fudan University biosafety level 3 (BSL-3) facility. The overall study was reviewed and approved by the SHAPHC Ethics Committee (approval no. 2020-Y008-01).

### Cell lines and Viruses

Vero E6, A549-Spike (A549 expressing SARS-CoV-2 S protein), and A549-ACE2 (A549 expressing human ACE2) cell lines were supplied by Shanghai Public Health Clinical Center, Fudan University. Pseudovirus of SAR2-CoV-2 was generated by Shanghai Public Health Clinical Center, and Fudan University and SARS-CoV-2-SH01 was from BSL-3 of Fudan University.

### B cell sorting and single cell RT-PCR

Samples of peripheral blood for serum or mononuclear cells (PBMCs) isolation were obtained from Shanghai Public Health Clinical Center, per 5 mL blood. PBMCs were purified using the gradient centrifugation method with Ficoll and cryopreserved in 90% heat-inactivated fetal bovine serum (FBS) supplemented with 10% dimethylsulfoxide (DMSO), storage in liquid nitrogen.

The fluorescently labeled S1 bait was previously prepared by incubating 5 μg of His tag-S1 protein with Anti His tag antibody-PE for at least 1 hr at 4 °C in the dark. PBMCs were stained using 7AAD, anti-human CD19 (APC), IgM [PE-Cy7], IgG (fluorescein isothiocyanate, FITC), PE labeled Antigen. Single antigen specific memory B cells were sorted on BD FACS Aria II into 96-well PCR plates (Axygen) containing 10 μl per well of lysis buffer (10 mM DPBS, 4 U Mouse RNase Inhibitor, NEB). Plates were immediately frozen on dry ice and stored at 80 C or processed for cDNA synthesis.

Reverse transcription and subsequent PCR amplification of heavy and light chain variable genes were performed using SuperScript III (Life Technologies). First and second PCR reactions were performed in 50 ml volume with 5ul of reaction product using PCR mixture (SMART-Lifesciences). PCR products were then purified using DNA FragSelect XP Magnetic Beads (SMART-Lifesciences) and cloned into human IgG1, lambda or kappa expression plasmids for antibody expression by seamless cloning method (see below). After transformation, individual colonies were picked for sequencing and characterization. Sequences were analyzed using IMGT/ V-QUEST (http://www.imgt.org/IMGT_vquest) and IgBlast (IgBLAST, http://www.ncbi.nlm.nih.gov/igblast).

### Expression and purification of human monoclonal antibodies

The antibody VH/VL and constant region genes were then amplified and cloned into expression vector pcDNA3.4 using SMART Assembly Cloning Kit (SMART-Lifesciences), subsequently antibodies plasmids were amplified in competent cells (SMART-Lifesciences). HEK293E cells were transfected using polyethylenimine (PEI, Sigma), after 4-5 days of cell culture, antibodies purification was processed from supernatants. Antibodies were purified by Protein A magnetic beads (SMART-Lifesciences) for 30 min at room temperature, then eluted by 100 mM glycine pH 3.0 and quickly neutralized by TrisCl pH 7.4.

### ELISA analyses

96-well plates (Falcon and MATRIX) were coated overnight at 4 °C with 0.5 μg/mL SARS-CoV-2 RBD-mFC (Novoprotein Scientific Inc.), and 0.6 μg/mL SARS-CoV-2 S-ECD (GenScript). After washes with PBST (SMART-Lifesciences), the plates were blocked using 3% non-fat milk in PBST for 1 h at 37°C. Series dilutions of antibodies in PBST were added to each well and incubated at 37°C for 1 h. Then the antbodies were removed and washed 3 times by PBST, HRP-conjugated anti-human IgG Fab antibody (Sigma) was added at the dilution of 1:10,000 in PBST containing 3% BSA (Sangon Biotech) and incubated at 37°C for 0.5 h. After washing with PBS/T three times, TMB solution (SMART-Lifesciences) was added to the microplate and incubated at room temperature for 5-10 min, followed by adding 1M HCl to terminate the reaction. The OD450 absorbance was detected by Synergy HT Microplate Reader (Bio-Tek). The curves and EC50 were analyzed by GraphPad Prism 8.0.

### Bio-layer Interferometry assay (BLI)

Measuring Kinetics of antibodies interact with S protein RBD using by Bio-layer Interferometry. RBD diluted in a 2-fold series by assay buffer (20mM MOPs, 50 nM KCl, pH 7.4).

### Flow cytometry assays

For Figure 4, flow cytometry analyses were performed to detect the binding abilities of antibodies to Spike protein in HEK293T cells freshly expressing of SARS-CoV-2, SARS-CoV and MERS-CoV. Briefly, 10 thousand cells in 100 μl were incubated with indicated antibodies for 30 min at room temperature, after twice washes then PE-labeled goat anti-human IgG-Fc antibody was added (1:5,000; Abcam) for 30 min, followed by flow cytometry analyses.

Flow cytometry analysis was also performed to detect the ACE2 competition (Figure 3C and Table S3). Then 10 thousand S expressing cells in 100 μl were first incubated with free ACE2 at 50nM for 30 min at room temperature, then different concentrations of antibodies were added for 30 min, followed by incubation with PE-labeled goat anti-human IgG-Fc antibody (1:5000; Abcam) for 30 min and analyzed by flow cytometry.

For Figure S3, to test S protein expression on cell membrane, recombinant ACE2-Cter-6XHis labeled by rabbit anti-His-PE antibody in 1.2:1 (*n:n*) ratio was added to 10 thousand cells expressing S protein. Detection was done similarly as above.

### Virus neutralization assay (pseudotyped and authentic)

Coding fragments of SARS-CoV-2 (YP_009724390.1), SARS-CoV (NP_828851.1) and MERS (AFS88936.1) Spike proteins were synthesized (GenScript) and cloned into pcDNA3.1. SARS-CoV-2, SARS-CoV and MERS pseudotyped viruses were produced as previously described^33^. Briefly, pseudovirus were generated by co-transfection of 293T cells with pNL4-3.Luc.R-E-backbone and the SARS-CoV-2 spike protein expression plasmid and the supernatants were harvested after 48 hr, and followed by centrifuge at 2,000 rpm for 5 min and stored in −80 °C.

The neutralization assay was performed as the following steps. Pseudovirus was diluted in complete DMEM mixed with or without an equal volume (50 μl) of diluted serum or antibody and then incubated at 37 °C for 1 h. The mixtures were then transferred to 96-well plate seeded with 20,000 293T-ACE2 cells for 12h and incubated at 37 °C for additional 48 hr. Assays were developed with bright glo luciferase assay system (Promega), and the relative light units (RLU) were read on a luminometer (Promega GloMax 96). The titers of neutralizing antibodies and activities of plasma were calculated as 50 % inhibitory dose (ID50) compared to virus control.

All experiments related to authentic virus were done in BSL-3. Monoclonal antibodies were incubated with 200 PFU SARS-CoV-2 SH01 at 37 °C for 1 hour before added into VERO E6 cell culture (96-well plate, 4 x 10^4^ cells per well), and the cells were continued for 48 hr before microscopic analyses for CPE (cytopathic effect).

## Supplementary Figure legends

**Tab.S1.**
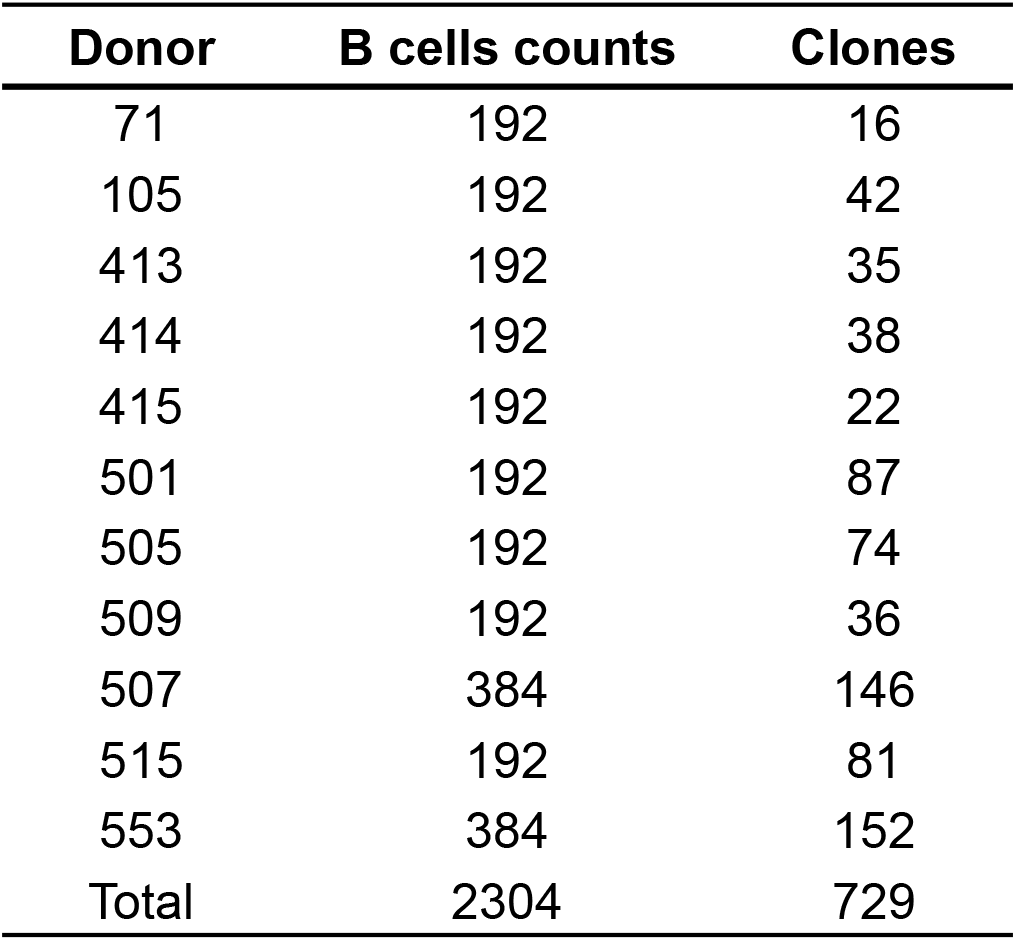
Related to Figure 1 and 2. Summary of numbers of obtained B cells and antibody clones from 11 convalescent patients.

**Tab.S2.** Related to Figure 1. Summary of the characteristics and symptoms of the 11 COVID-19 patients.

| Patient characteristics |  |  |  |  |  |  |  |  |  |  |  |
| --- | --- | --- | --- | --- | --- | --- | --- | --- | --- | --- | --- |
| Patient number | 71 | 105 | 413 | 414 | 415 | 501 | 505 | 507 | 509 | 515 | 553 |
| Gender | Female | Male | Female | Male | Female | Male | Female | Male | Male | Female | Male |
| Age(years) | 35 | 30 | 48 | 56 | 65 | 29 | 60 | 15 | 54 | 59 | 40 |
| Sampling time point(after hospitalization) | 10 days | 18 days | 19 days | 16 days | 16 days | 12 days | 15 days | 9 days | 1 days | 9 days | 25 days |
| Fever | Yes | Yes | Yes | Yes | No data | Yes | Yes | Yes | No data | Yes | Yes(with myalgia) |
| Other infomations | Traveled in Shanghai from Wuhan at Jan 21st | Jan 13rd to 17th, spent time with a confirmed patient | Been to Wuhan at Jan 15th | Passed by Wuhan at Feb 23rd | Spent time with a low fever friend at Jan 19th | No data | Passing by Wuhan at Jan 17th for 1 hour | No data | Been close with a confirmed patient at Jan 14th | No data | Been to Wuhan for working at Jan 11st |

**Tab.S3.** Related to Figure 2 and 3. Summary of the performance of the top 30 monoclonal antibodies in the indicated assays.

| Antibodies | Neutralization<br>of Authentic<br>IC50 (nM) | Neutralization<br>of Pseudoviral<br>IC50 (nM) | RBD+S-ECD<br>Binding ELISA<br>Kd (nM) | Blocking<br>ELISA IC50<br>(nM) | FCA<br>S protein<br>Binding<br>(Positive %) | FCA<br>Competition<br>to ACE2 (50nM) | RBD ELISA<br>Binding<br>(OD450) | S-ECD ELISA<br>Binding<br>(OD450) |  |
| --- | --- | --- | --- | --- | --- | --- | --- | --- | --- |
| **515-1** | **~ 6** | **26.11** | **0.057** | **0.821** | **53.8** | **+** | **3.296** | **1.99** |  |
| **505-5** | **~ 6** | **26.37** | **0.059** | **0.175** | **56.63** | **++** | **3.357** | **2.044** |  |
| **413-2** | **~ 100** | **>100** | **0.06** | **non** | **39.32** | **+++** | **0.163** | **2.733** |  |
| 414-2 | non | 12.721 | 0.074 | non | 64.16 | + | 4.044 | 2.994 |  |
| **553-49** | **~ 10** | **>100** | **0.076** | **non** | **32.561** | **++** | **2.572** | **1.908** |  |
| **505-3** | **~ 3** | **2.313** | **0.08** | **0.719** | **61.64** | **++** | **3.185** | **2.107** |  |
| **553-15** | **~ 30** | **3.628** | **0.089** | **0.325** | **41.68** | **+** | **3.359** | **2.458** |  |
| 105-38 | non | 24.44 | 0.099 | 0.122 | 33.11 | + | 3.735 | 2.32 |  |
| 415-5 | non | >100 | 0.148 | 12.03 | 44.53 | ++ | 3.777 | 2.308 |  |
| 553-20 | non | >100 | 0.209 | non | 63.45 | non | 1.83 | 0.249 |  |
| 105-9 | non | >100 | 0.257 | 4.663 | 56.26 | + | 2.641 | 1.095 |  |
| **553-63** | **~ 6** | **3.569** | **0.302** | **0.237** | **73.73** | **+++** | **3.464** | **2.114** |  |
| **414-1** | **1.75** | **3.085** | **0.31** | **2.964** | **58.53** | **+** | **3.772** | **2.034** |  |
| 553-13 | non | non | 0.683 | non | 77 | +++ | 0.149 | 2.708 |  |
| 415-6 | non | non | 0.701 | non | 17.58 | ++ | 3.895 | 2.712 |  |
| 553-27 | non | non | 0.769 | non | 41.63 | +++ | 3.409 | 3.081 |  |
| 553-18 | non | non | 0.853 | non | 61.04 | +++ | 0.181 | 0.302 |  |
| 413-3 | non | non | 2.655 | non | N.D. | ++ | 1.82 | 0.924 |  |
| 105-43 | non | non | 3.042 | non | 34.58 | + | 2.999 | 1.101 |  |
| 414-4 | non | non | 3.119 | non | N.D. | non | 2.886 | 0.77 |  |
| 105-28 | non | non | 4.244 | non | N.D. | non | Not tested | Not tested |  |
| 105-41 | non | non | 4.811 | non | N.D. | non | Not tested | Not tested |  |
| **553-60** | **~ 10** | **3.877** | **5.753** | **non** | **36.75** | **++** | **2.996** | **1.4** |  |
| 414-3-2 | non | non | 6.65 | non | 30.68 | ++ | 3.515 | 2.428 |  |
| 553-17 | non | non | 17.31 | non | 42.77 | + | 2.288 | 0.586 |  |
| **505-8** | **~ 4** | **4.186** | **17.62** | **non** | **30.44** | **+++** | **0.181** | **0.248** |  |
| 105-25 | non | non | 55.15 | non | N.D. | non | Not tested | Not tested |  |
| 414-3-1 | non | non | 64.63 | non | 17.96 | non | 2.754 | 1.229 | +++ : 500-folds competition to ACE2 |
| 505-17 | non | non | 78.63 | non | 29.76 | + | 0.135 | 0.141 | ++ : 50-folds competition to ACE2 |
| **515-5** | **~ 100** | **4.943** | **119.7** | **non** | **34.08** | **+** | **0.227** | **0.263** | + : 5-folds competition to ACE2 |
|  |  |  |  |  |  |  |  |  | N.D. : Non-Detectable |

**Fig. S1.**
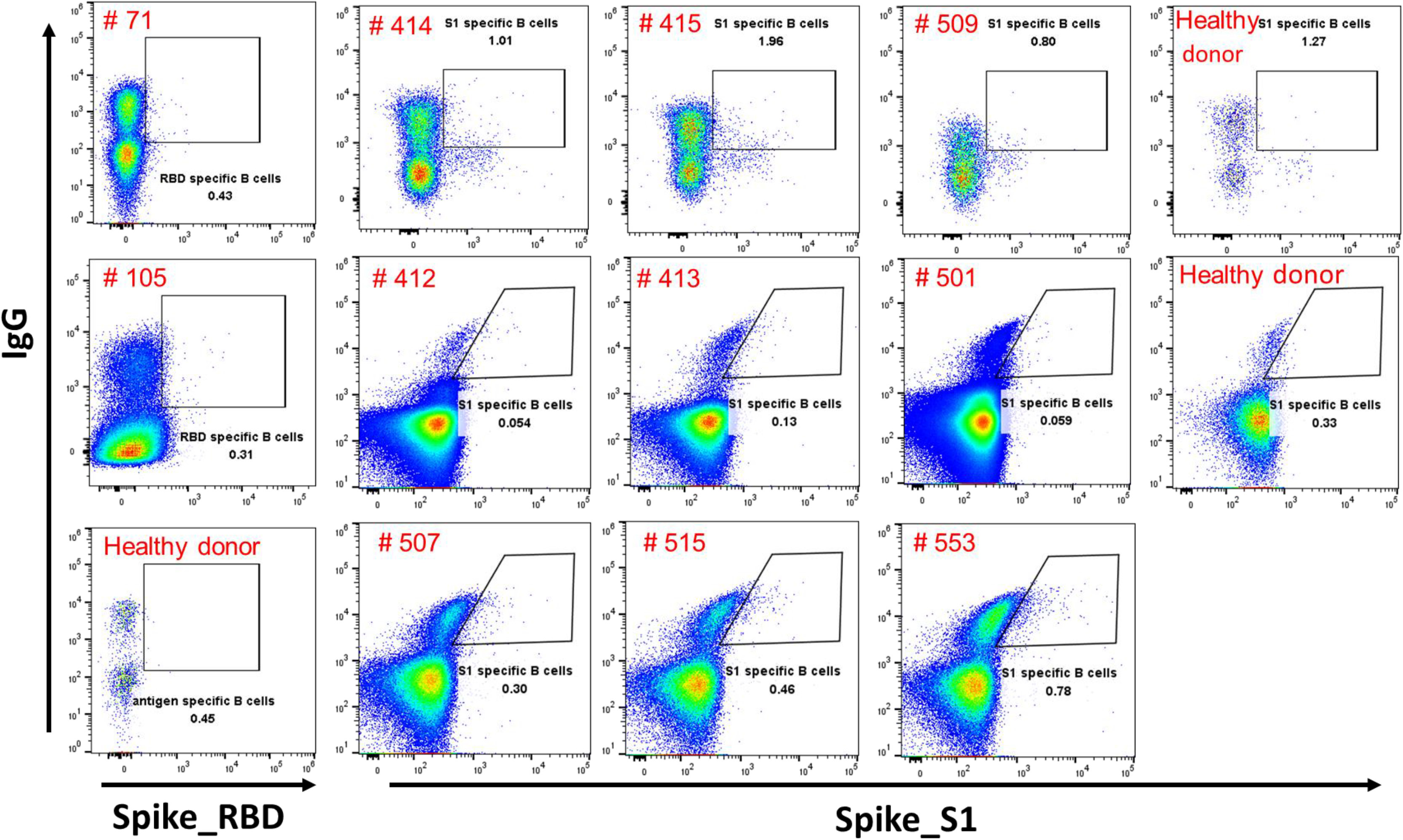
Related to Figure 1. Flow cytometry B cell sorting from PBMCs of 11 convalescent patients.

**Fig. S2.**
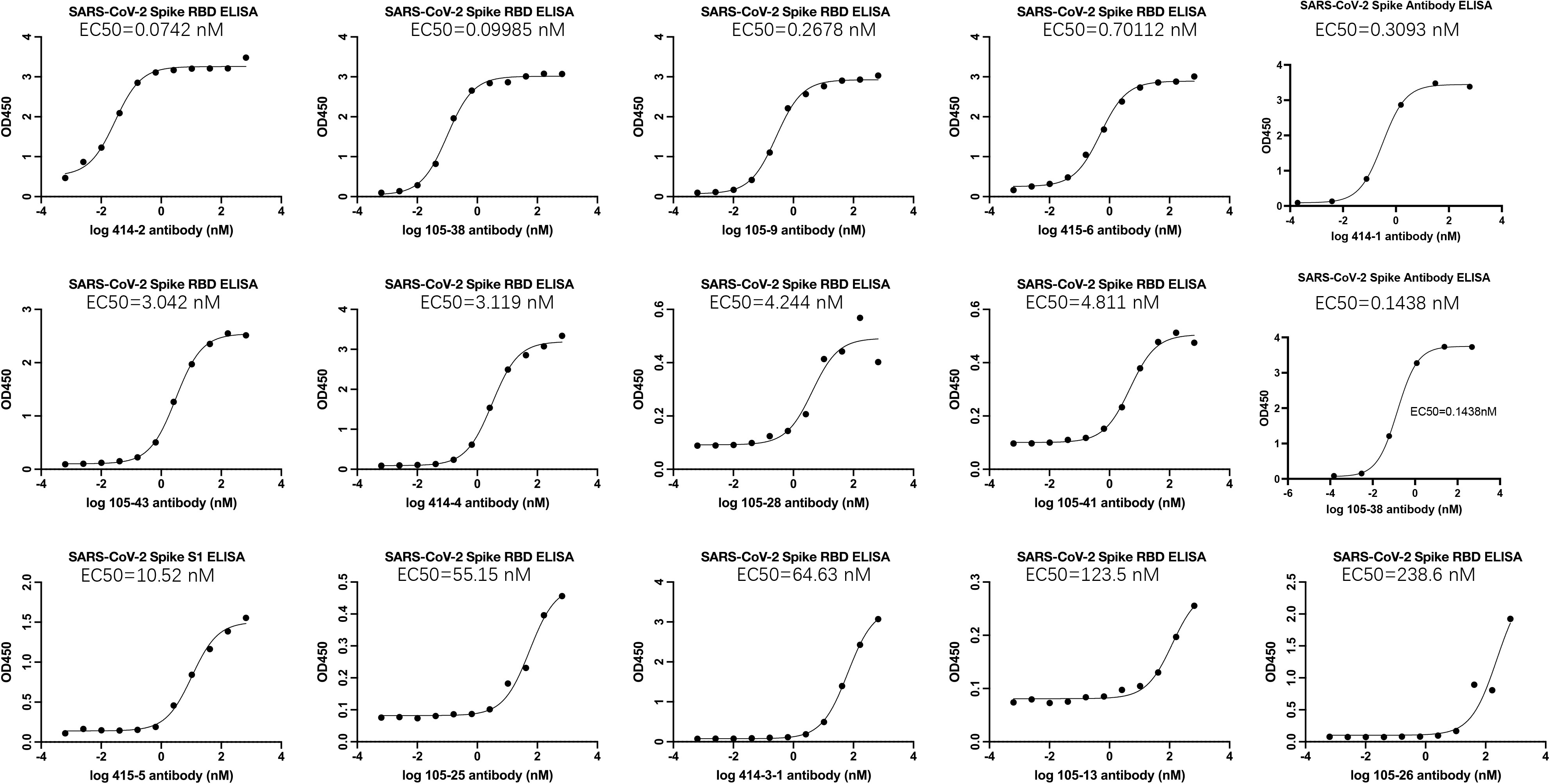
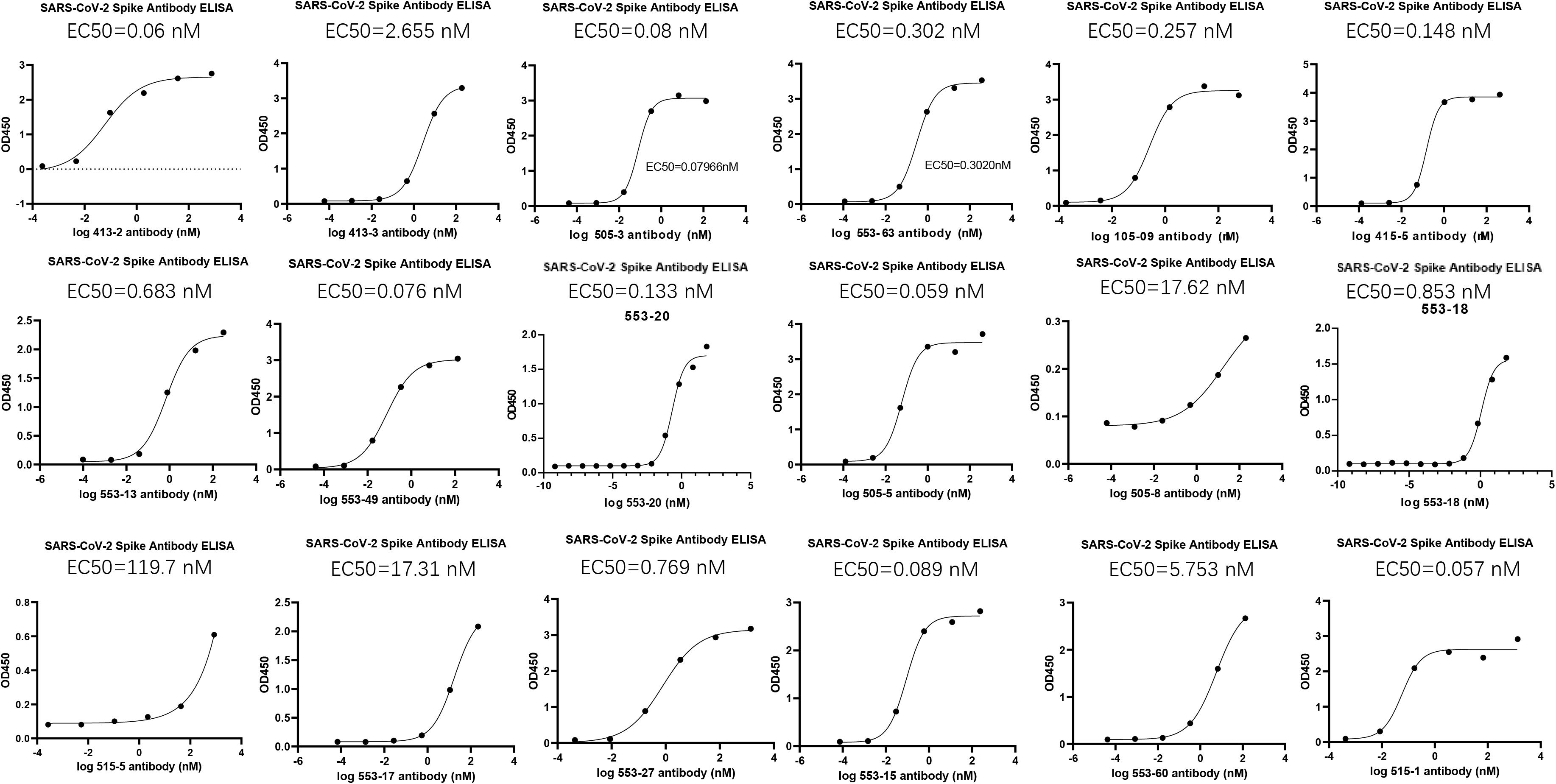
Related to Figure 2. Summary of ELISA binding data of the select 33 SARS-CoV-2 specific monoclonal antibodies.

**Fig. S3.**
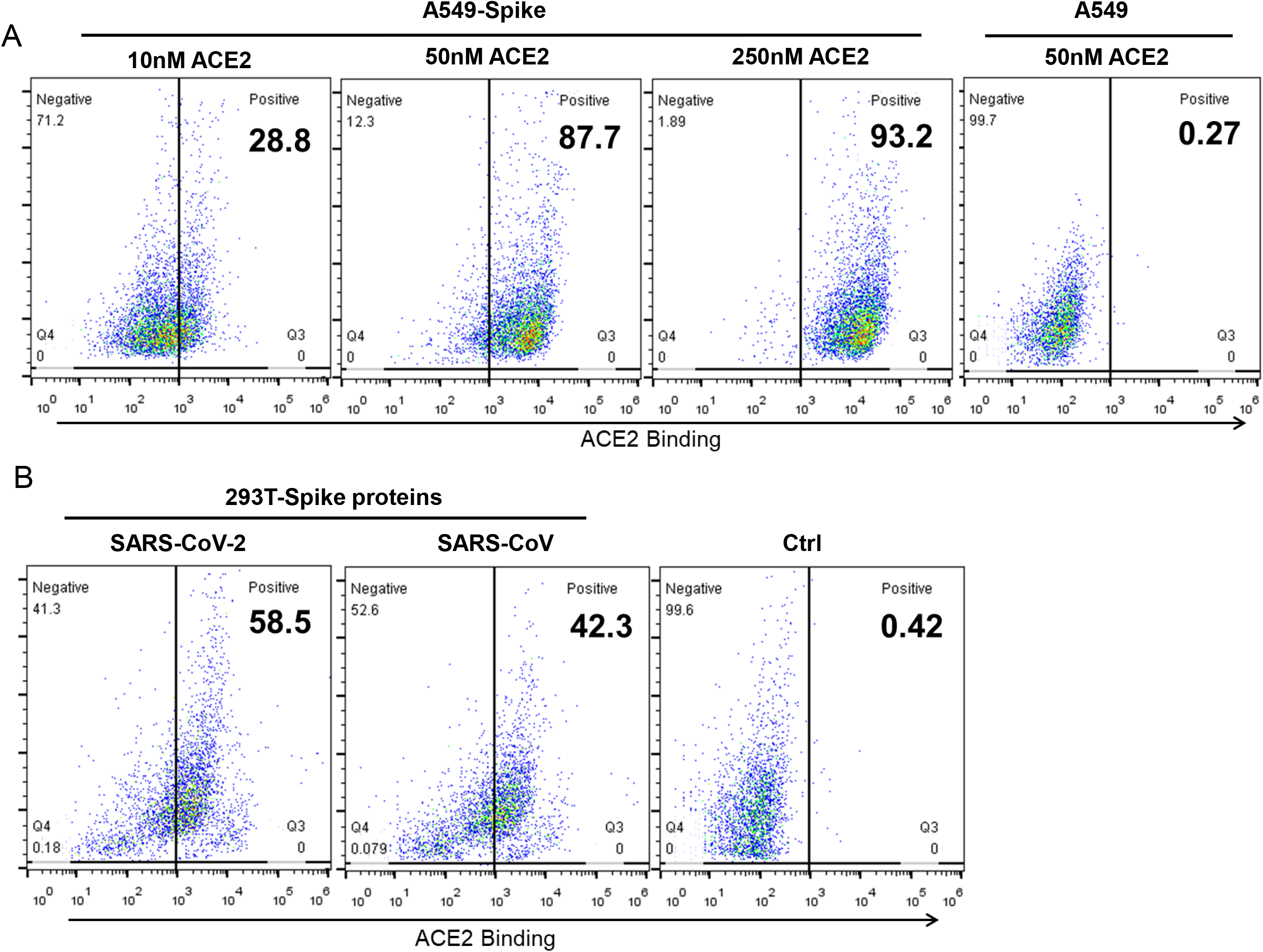
Related to Figure 3 and 4. ACE2 binding to S protein expressed on A549 and HEK293T cell membrane. (A) ACE2 binding to S protein expressed in A549 detected by flow cytometry. (B) ACE2 binding to S proteins from SARS-CoV-2 and SARS-CoV expressed on HEK293T membrane. Non-transfected A549 and HEK293T were used as controls.

**Fig. S4.**
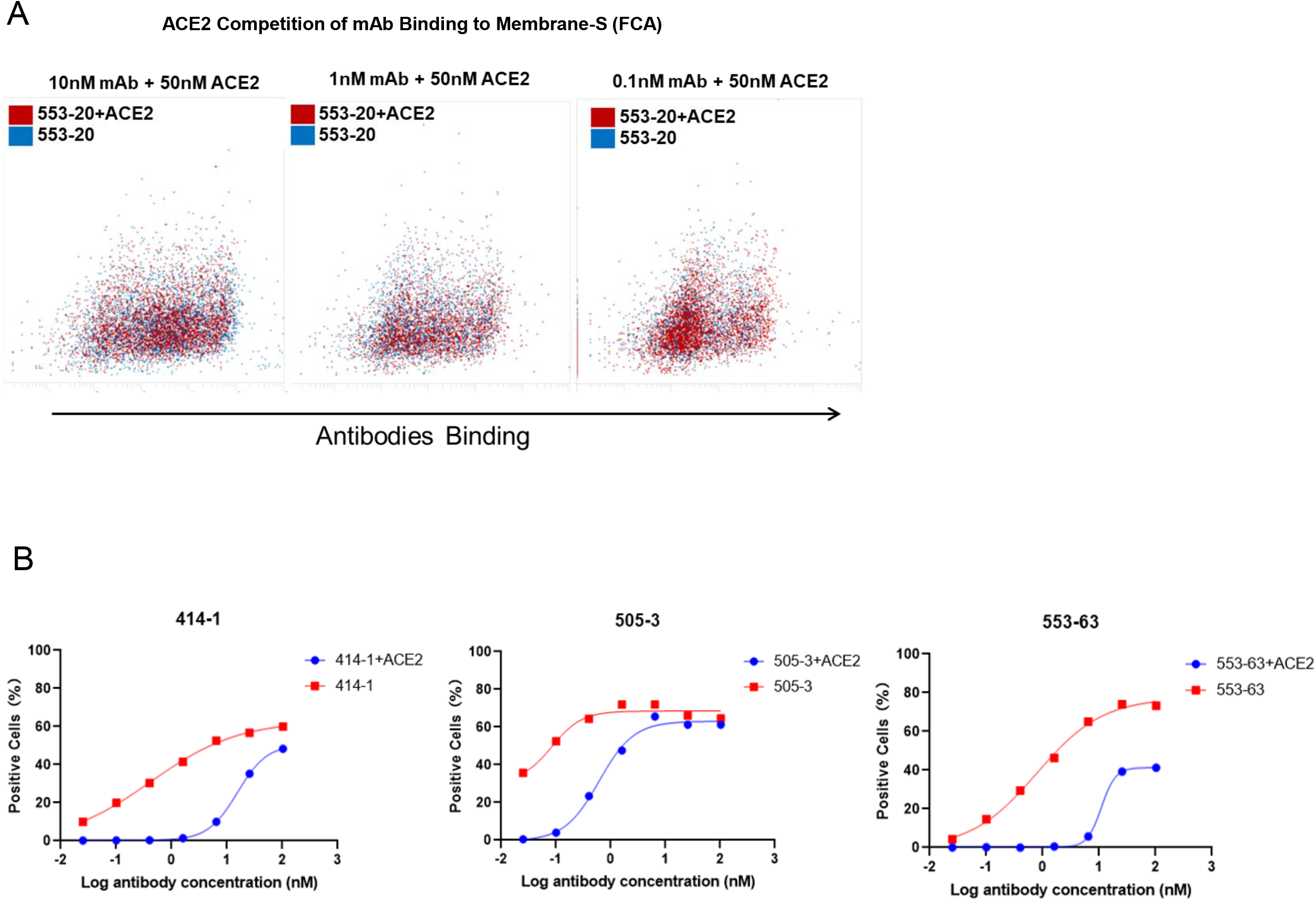
S4 Related to Figure 3. (A-B) ACE2 competition assay of 553-20, 414-1, 505-3 and 553-63 using flow cytometry.

